# Time course of alterations in adult spinal motoneuron properties in the SOD1(G93A) mouse model of ALS

**DOI:** 10.1101/2020.05.19.105007

**Authors:** Seoan Huh, Charles J. Heckman, Marin Manuel

**Author notes:** **Corresponding author**: Marin MANUEL, SPPIN - Saints-Pères Paris Institute for the Neurosciences, Université de Paris, 45, rue des Saints-Pères, 75006 Paris, France. **Author contribution**: Conceptualization: S.H., C.J.H., and M.M.; Methodology: S.H., C.J.H., and M.M.; Investigation: S.H.; Formal analysis: S.H. and M.M.; Software: M.M.; Visualization: S.H. and M.M.; Writing - Original Draft: S.H.; Writing - Review & Editing: S.H., C.J.H., and M.M.; Funding Acquisition: C.J.H and M.M.; S.H., C.J.H., and M.M. approved the final version of this manuscript.

## Abstract

Although ALS is an adult-onset neurodegenerative disease, motoneuron electrical properties are already altered during embryonic development. Motoneurons must therefore exhibit a remarkable capacity for homeostatic regulation to maintain a normal motor output for most of the life of the patient. In the present paper, we demonstrate how maintaining homeostasis could come at a very high cost. We studied the excitability of spinal motoneurons from young adult SOD1(G93A) mice to end-stage. Initially homeostasis is highly successful in maintaining their overall excitability. This initial success, however, is achieved by pushing some cells far above the normal range of passive and active conductances. As the disease progresses, both passive and active conductances shrink below normal values in the surviving cells. This shrinkage may thus promote survival, implying the previously large values contribute to degeneration. These results support the hypothesis that motoneuronal homeostasis may be “hyper-vigilant” in ALS and a source of accumulating stress.

**Significance Statement:** During ALS, motoneurons exhibit a remarkable ability to maintain a normal motor output despite continuous alterations of their electrophysiological properties, up to the point when overt symptoms become apparent. We show that this homeostatic process can sometimes push motoneurons beyond the normal range, which may be causing long-lasting harm.

## Introduction

ALS is a fatal neurodegenerative disorder characterized by progressive loss of cortical and spinal motoneurons. Five to ten percent of the ALS cases are familial (FALS), and ~20% of the FALS cases are due to mutations in the Superoxide Dismutase1 (SOD1) gene (Rosen et al., 1993). Studies in the mutant SOD1 (mSOD1) mouse and other animal models have revealed that multiple cellular functions become impaired as the disease progresses (Ilieva et al., 2009; Kanning et al., 2009; Johnson and Heckman, 2010; Brownstone and Lancelin, 2018), implying that homeostatic mechanisms are failing. Yet, despite an early surge in motoneuron excitability in the embryonic state (Kuo et al., 2004; van Zundert et al., 2008; Pieri et al., 2009; Martin et al., 2013), mutant mice maintain a normal motor output for several months, suggesting the involvement of very potent and effective homeostatic process to maintain the intrinsic excitability of the motoneurons.

The standard measure of the net excitability of motoneurons is the relation between the frequency of firing and the amplitude of the injected current. Multiple ionic currents contribute to this frequency-current (F-I) function relationship. Chief among them are persistent inward currents (PICs), which are mediated by voltagegated Na+ and Ca2+ channels. In neonatal mSOD1 mice, PICs in spinal motoneurons become aberrantly large (Quinlan et al., 2011). On its own, this change would increase F-I gain, but it is compensated by commensurate increases in the leak currents that set the input conductance of the cell, so that the net excitability remains constant (Quinlan et al., 2011). These abnormal changes show that F-I homeostasis during the neonatal period is achieved via a compensatory mechanism. If, however, these compensatory increases persist into the young adult state, then the continued distortions in input conductances and PIC amplitudes could induce a substantial stress within the motoneurons. Increasing the number of leak conductances and voltage-gated channels implies a higher energy expenditure to maintain the resting membrane potential and ionic gradients across the membrane, which already occupies a large share of the metabolic budget of neurons (Attwell and Laughlin, 2001; Herculano-Houzel, 2011; Howarth et al., 2012). One must also consider the additional burden associated with the increased housekeeping tasks such as lipid synthesis, trafficking of organelles and protein synthesis, which account for 25–50% of the energy budget of neurons (Rolfe and Brown, 1997; Attwell and Laughlin, 2001). In this scenario, the initial success in the homeostatic regulation of net excitability would come at a severe cost, likely accelerating the onset of degeneration. Indeed, neurons function on a very restricted energy budget that is independent of their size (Herculano-Houzel, 2011). On the other hand, if homeostatic processes instead successfully return input conductance and PIC values to normal ranges before the onset of denervation, then homeostasis for the F-I function would reduce stress on the cell and effects on subsequent degeneration would likely be small.

Here, we investigated how the homeostatic processes required to maintain normal excitability (current onset and F-I gain) develop as the disease progresses. We undertook the first *in vivo* voltage-clamp studies of motoneurons in the SOD1(G93A) mouse model of ALS and pursued these measurements across a wide range of ages, from P30 to P120. This age range spans the young adult period, the onset of denervation period (~P50, Pun et al., 2006), and the development of overt symptoms (~P90). Because adult mSOD1 motoneurons tend to have a larger input conductance than controls (Delestrée et al., 2014), we hypothesized that input conductance and PIC values would continue to grow in the young adult state. Our results supported this hypothesis, revealing, in fact, a continual increase in the amplitude of these parameters, followed by a collapse, so that motoneurons surviving beyond the onset of overt symptoms (~P90) had aberrantly small values of each. These results are consistent with the possibility that homeostasis for excitability is not weak but excessively strong and that this overreaction contributes to subsequent degeneration (Mitchell and Lee, 2012).

## Methods

### Animals

This study was performed in strict accordance with the recommendations in the Guide for the Care and Use of Laboratory Animals of the National Institutes of Health. All of the animals were handled according to protocols approved by Northwestern University’s institutional animal care and use committee (IACUC). All surgery was performed under sodium pentobarbital anesthesia, and every effort was made to minimize suffering. Because of the reproducible and stereotyped progression of symptoms, we chose to use mice overexpressing the human SOD1(G93A) gene, in which glycine has been substituted by alanine at residue 93 as a mouse model of ALS. Hemizygous B6SJL.SOD1(G93A) transgenic mutant males were bread with B6SJL F1 females (obtained from the Jackson Laboratory, Bar Harbor, ME)(Leitner et al., 2009). Offspring were genotyped and the transgene copy number was compared to a housekeeping gene at Transnetyx (Cordova, TN). Only mice with relative copy number >45 were used in this study. The control group consisted of non-transgenic littermates, with the same B6SJL background (WT). 33 animals of either sex were used in this study. Because these experiments tend to have a higher success rate with larger animals, our sample was biased towards males (27 males and 6 females). Animals were divided into different age groups for analysis: animals whose age was <60 days-old were categorized as P30-60 (N=9 mice), animals whose age was ≥60 and <90 as P60-90 (N=12 mice), and animals older than ≥90 days-old were classified as P90-120 (N=12 mice).

### In vivo preparation

Procedures were similar to those in our previous studies (Manuel and Heckman, 2011). Initially, atropine (0.2 mg/kg) was administered subcutaneously to prevent salivation. Ten minutes later, anesthesia was initiated with an intraperitoneal injection of pentobarbital sodium (70 mg/kg) and maintained by intravenous infusion of supplemental doses of pentobarbital (6 mg/kg) mixed in perfusion solution containing 4% glucose, 1% NaHCO_3_, and 14% Plasmion. The trachea was cannulated, allowing the mouse to be artificially ventilated with 100% O_2_. The end-tidal PCO_2_ was continuously monitored and maintained around 4% by adjusting ventilator parameters including respiratory rate around 120-150 bpm and tidal volume around 0.09-0.23 mL. The hind limb muscles were then dissected; biceps femoris was removed to expose the sciatic nerve, which was thereafter placed over a stimulating electrode. A laminectomy was performed at the T13-L1 level and the L3-L4 spinal segments were exposed. To prevent the spinal cord from dehydration, a custom made bath was affixed using silicone elastomer and covered with mineral oil. To locate the motoneurons of interest, the sciatic nerve was stimulated at 1.8-2× the minimum intensity required to observe an afferent volley.

### Electrophysiology

Intracellular recordings of spinal motoneurons were performed by impaling them with glass micropipette electrodes filled with 3M KCl with a resistance of 8-15 MΩ. Motoneurons were identified by the presence of antidromic action potential from stimulation of the sciatic nerve. Cells with unstable resting membrane potential (RMP) or with RMP more depolarized than −50 mV were excluded from the analysis. We record a median of 3 cells per animal (average ± SD 3.1±1.7 cells per animal, mode 2 cells/animal, N=33).

PICs were recorded in discontinuous voltage-clamp mode, with switching rates of 6-8 kHz. Clamp feedback gain was between 0.3 and 1.5. In addition to the feedback gain from the Axoclamp amplifier, an additional low-frequency feedback loop with a gain of 11 and a cutoff of −3 dB at 0.3 kHz was used to improve voltage control in such large cells as motoneurons (Lee and Heckman, 1998). Monitoring outputs were observed at all times to assure reasonable settling of the electrode. To record PICs, a slow triangular voltage ramp (−80 to −40 mV) was applied (Lee and Heckman, 1998). Leak current was determined by fitting a regression line through the subthreshold region (−80 to −65 mV) of the I-V function. Then, this leak was subtracted from the total function to determine the PIC amplitude (measured both on the ascending and descending part of the ramp). In addition, the voltage at which the PIC was maximal on each part of the ramp was recorded (PIC peak voltage). The PIC onset voltage was estimated on the leak-subtracted trace as the point where the curve started to visibly deviate downward from the horizontal. Input conductance was estimated as the slope of the leak current.

To assess the intrinsic properties of the motoneuron in current-clamp mode, we measured the frequency-current (F-I) relationship of action potential firing. The F-I relationship was determined based on the firing produced by a triangular current injection. The interspike frequency was then plotted against the intensity of the injected current. All these measurements were carried out in the discontinuous current-clamp mode of the Axoclamp 2A amplifier, with switching rates of 6-8 kHz. Mouse motoneurons possess two regimes of firing: a sub-primary range of firing, followed by a linear primary range (Manuel et al., 2009; Manuel and Heckman, 2011). The primary range was identified visually starting from the top of the ramp and going backward towards the beginning (for the ascending ramp) of forward towards the end (for the descending ramp). A linear portion of the instantaneous frequency, with low variability, can generally be easily identified before a sudden change of slope or an increase in firing variability. The “gain” of the F-I relationship was determined by fitting a regression line over the linear range so identified (separately on the ascending and descending phase of the ramp). The other parameters measured in the F-I relationship were: the current at which the first action potential (AP) fires on the ascending ramp (recruitment current). the current at which the last AP fires on the descending ramp (current at de-recruitment); the voltage threshold for spiking, which was determined as the voltage where the slope of the membrane voltage reaches 10 mV/ms before the first AP (Sekerli et al., 2004); the current and firing frequency at the transition between the sub-primary and primary ranges (Manuel and Heckman, 2011).

### Data analysis

Recordings were acquired and analyzed using Spike2 v.7 (CED, Cambridge, UK). Data were analyzed using the scientific python (v.3.7.4) ecosystem: Pandas v.1.0.5 (McKinney, 2011), SciPy v.1.5.2 (Virtanen et al., 2020), DABEST v.0.3.0 (Ho et al., 2019), statsmodels v.0.12.0 (Seabold and Perktold, 2010) and scikit-learn v.0.23.0 (Pedregosa et al., 2011). Figures were generated using matplotlib v.3.1.3 (Hunter, 2007) and seaborn v.0.10.0 (Waskom et al., 2020).

### Statistical analysis

All data are reported as mean ± standard deviation with their respective sample size. Each cell is treated as an independent observation, and the reported N refers to the number of cells, unless otherwise specified. No test was performed to detect outliers, and no data points were excluded from the analysis. When comparing between WT and mSOD1 samples, we focus on estimation statistics that rely on effect sizes and confidence intervals (95%CI), rather than null hypothesis significance testing, as recommended by several scientific societies and editorial boards (Bernard, 2019; Makin and Orban de Xivry, 2019; Wasserstein et al., 2019; Michel et al., 2020). Unless otherwise specified, effect sizes are reported as differences of means and Hedges’ *g* (Hedges, 2016). Where appropriate, data are presented as Cumming plots (Cumming, 2012) generated using DABEST. In these plots, the raw data are plotted as swarms of points. In addition, the mean ± standard deviation of each group is plotted as a notched line immediately to the right of each group. The effect size and bootstrapped 95% CIs are plotted on separate axes beneath the raw data. Confidence intervals were bias-corrected and accelerated, and are displayed with the bootstrap distribution of the mean; resampling was performed 5000 times (Ho et al., 2019). Welch’s *t*-test (Welch, 1947) results are provided for information only. ANCOVA was performed using statsmodels’ OLS routines, fitting the model “PIC amplitude ~ input conductance * Genotype”. No significant interaction term was detected for any of the age groups considered, and the model was run again without interaction. Dimensionality reduction was performed using scikit-learn’s PCA. The 21 electrophysiological features (which did not include genotype or age) of our dataset were centered and scaled then projected on a 5D space. For analysis, only the first three principal components were considered.

## Results

We set out to compare neuronal excitability, in 103 motoneurons from SOD1(G93A) mice (mSOD1; 53 motoneurons), and their non-transgenic (WT; 50 motoneurons) littermate, aged between 31 and 123 days old. Neuronal excitability was estimated on the response of the motoneurons to a triangular ramp of current in current-clamp mode. In addition, PICs were measured in each motoneuron using a triangular voltage ramp in voltage-clamp mode. Most of the values measured on the ascending and descending phases of the ramps, both in voltage-clamp and current-clamp were very strongly correlated (PIC amplitude on ascending and descending ramps: *r^2^*=0.75; PIC onset voltage: *r^2^*=0.89; PIC peak voltage: *r^2^*=0.90; recruitment and de-recruitment currents: *r^2^*=0.90). For this reason, we will mostly focus on the parameters measured on the ascending ramp.

Given the large period considered here, one also needs to consider whether alterations in motoneuron properties were direct effects of the mutation or merely shifts in the distribution of the properties caused by the progressive loss of a fraction of the motoneurons, starting with the least excitable FF motoneurons (Pun et al., 2006; Hegedus et al., 2007, 2008; Martínez-Silva et al., 2018). For this reason, we have split our dataset into three age groups. First, the young adult group (P30-60) corresponds to pre-symptomatic animals, with little to no neuromuscular-junction denervation (denervation starts at >P50 in FF motor units (Pun et al., 2006)). The presymptomatic group (P60-90) corresponds to a group where there are no overt motor symptoms despite a substantial loss of distal axons (Pun et al., 2006) and the beginning of cell death in the spinal cord (Kanning et al., 2009; Lalancette-Hebert et al., 2016). The symptomatic group (P90-120) contains animals showing overt motor symptoms, and which have lost a significant proportion of their motoneurons, particularly in the FF and FR population. Because motoneurons are still intact in the young adult group, changes in electrical properties can be directly interpreted as the direct result of the mutation, rather than the result of a shift in the population average caused by the loss of a fraction of the motoneurons. This age group will therefore be the main focus of the rest of our analysis.

### Motoneurons from young adult mutant mice have overly large PICs

In motoneurons, one of the major determinants of excitability are PICs, which were measured in response to a slow (5 mV/s) ramp in voltage-clamp mode. The amplitude of the PIC was measured on the leak-subtracted trace at the point where the downward deflection of the trace was maximal (see Figure 1A-B). We also measured the PIC activation voltage, and the voltage at which the PICs reached their peak.

**Figure 1:**
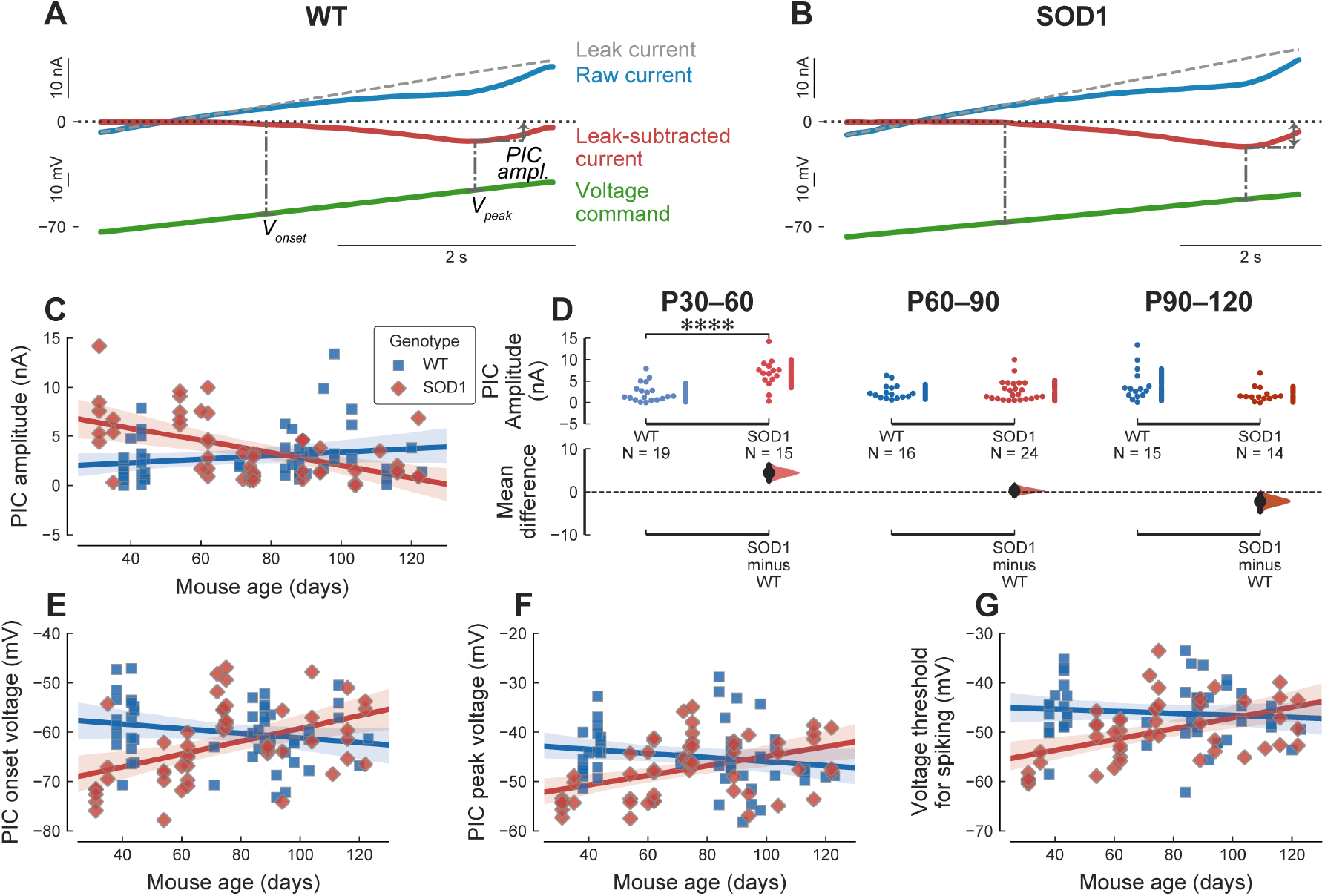
PIC amplitude is larger is young adult mSOD1 mice. **A**. Example of a PIC recording from a P43 WT mouse. The green bottom trace is the ascending part of the voltage ramp. The top blue trace is the raw current. The dashed line shows the leak current estimated by fitting a straight line in the subthreshold potential region, and which is used to measure the input conductance of the cell. The leak-subtracted current trace is obtained by subtracting the leak current from the raw current trace. The dash-dotted lines show some of the measurements: *PIC amp*.: PIC amplitude, measured at the point of largest deflection on the leak-subtracted trace; *V_onset_*: voltage at which the PICs start to activate; *V_peak_*: voltage at which PICs reach their maximum. **B**. Example of a PIC recording from a P35 mSOD1 mouse. Same organization as in A. **C**. Plot of the amplitude of the PICs (in nA) vs. age in WT (blue square) and mSOD1 mice (red diamonds). The solid lines correspond to the linear regression lines with 95% confidence intervals (shaded areas). WT: slope=0.12 nA/week 95%CI[−0.068-0.32], *r^2^*=0.034 (*p*=0.2); SOD1: slope=−0.45 nA/week 95%CI[−0.64–−0.25], *r^2^*=0.29 (*p*=3.4e-05). **D**. Breakdown of the difference in PIC amplitude between WT and mSOD1 animals by age groups. P30-60 WT: 2.30±2.17 nA, N=19 vs. mSOD1: 6.73±3.28 nA, N=15; *g*=1.59 95%CI[0.63-2.45]; *t*(32)=−4.50, *p*=0.00016. P60-90 WT: 2.47±1.69 nA, N=16 vs. mSOD1: 2.75±2.40 nA, N=24; *g*=0.12 95%CI[−0.53-0.65]; *t*(38)=−0.42, *p*=0.67. P90-120 WT: 4.12±3.69 nA, N=15 vs. mSOD1: 1.90±1.78 nA, N=14; *g*=−0.74 95%CI[−1.26-0.01]; *t*(27)=2.08, *p*=0.05. **E**. Evolution of the membrane potential at which the PICs start to activate (PIC onset voltage) vs. age. WT: slope=−0.33 mV/week 95%CI[−0.77-0.11], *r^2^*=0.045 (*p*=0.14). SOD1: slope=0.91 mV/week 95%CI[0.41-1.4], *r^2^*=0.21 (*p*=0.00063). **F**. Evolution of the membrane potential at which the PICs reach their peak (PIC peak voltage) vs. age. WT: slope=−0.29 mV/week 95%CI[−0.73-0.16], *r^2^*=0.034 (*p*=0.2). SOD1: slope=0.68 mV/week 95%CI[0.28-1.1], *r^2^*=0.18 (*p*=0.0014). **G**. Evolution of the voltage threshold for spiking (measure in current-clamp mode) vs. age. WT: slope=−0.14 mV/week 95%CI[−0.57-0.28], *r^2^*=0.0096 (*p*=0.5). SOD1: slope=0.77 mV/week 95%CI[0.34-1.2], *r^2^*=0.23 (*p*=0.0007).

In WT animals, PIC amplitude remained roughly steady during the age period studied (Figure 1C). However, in mSOD1 motoneurons, PIC amplitudes started higher than in WT animals at the youngest stages, then decreased strikingly over time (Figure 1C). When broken down by age groups (Figure 1D), our results show that, in the young adult stage (P30-60), the SOD1 mutation lead to an almost 3× increase in PIC amplitudes compared to WT motoneurons. This increase appears to be a continuation of the trend observed in neonates, where PICs were already increased ~2 fold compared to controls (Quinlan et al., 2011). In the presymptomatic group (P60-90), however, mSOD1 PICs shrunk to the same amplitude as WT motoneurons. At symptomatic stages (P90-120), the trend seen in young animals is reversed: PICs are much smaller in mSOD1 motoneurons compared to WT motoneurons (Table 1).

**Table 1:**
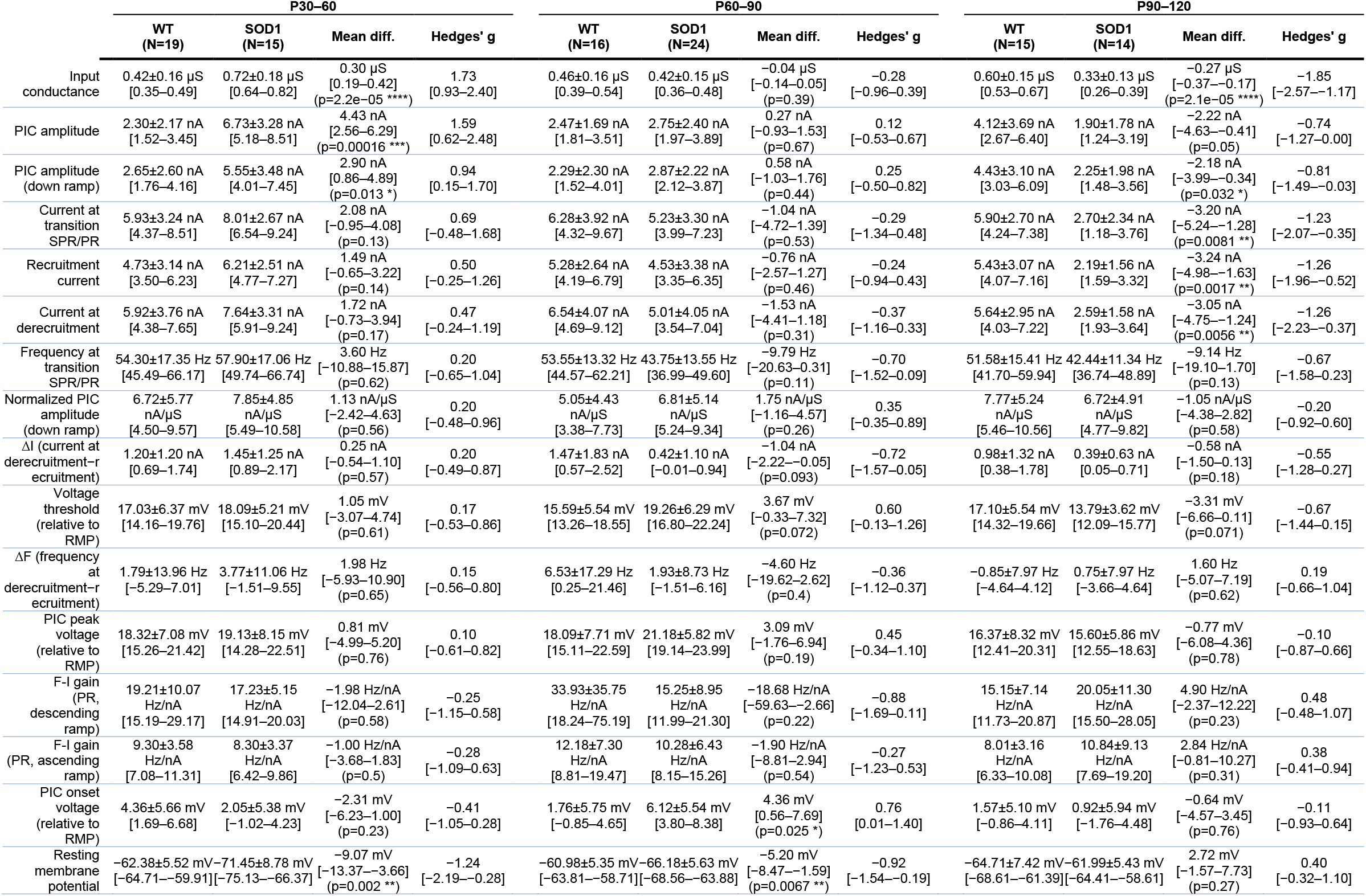
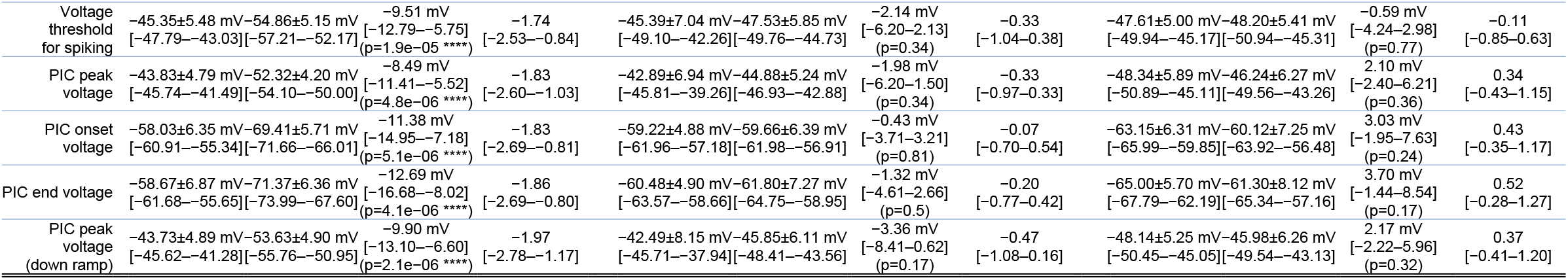
summary of the properties of the motoneurons, broken down by age range. The table shows the various properties that were measured in each motoneuron. The rows are sorted in descending order of effect size at P30-60. WT and mSOD1 columns show mean values ± standard deviation, with the 95%CI around the mean below. The “Mean diff.” column shows the mean difference between mSOD1 and WT population, with 95% CI and the result of Welch’s test (for information). The “Hedges’ *g*” column shows the effect size and its 95%CI below.

In addition to its effect on PIC amplitude, SOD1 mutation also affected the voltage at which PICs are recruited. The “PIC onset voltage”, measured as the voltage at the point where the leak-subtracted motoneuron I-V curve initially begins to curve downward, was hyperpolarized by 10 mV in young mSOD1 mice compared to WT controls (WT: −58.03±6.35 mV, N=19 vs. mSOD1: −69.41±5.71 mV, N=15; *g*=−1.83 95%CI[−2.64–−0.78]; *t*(32)=5.49, *p*=5.1e-06), and then slowly increased over time to match the value in WT motoneurons by endstage (Figure 1E). The SOD1 mutation had a similarly large effect on the voltage at which the PIC reached its maximum in young animals (WT: −43.83±4.79 mV, N=19 vs. SOD1: −52.32±4.20 mV, N=15; *g*=−1.83 95%CI[−2.63–−1.04]; *t*(32)=5.50, *p*=4.8e-06). The peak voltage then increased progressively over time, paralleling the onset voltage (Figure 1F).

These profound changes in PIC amplitude and activation voltage would be expected to have an impact on the firing properties of motoneurons. We, therefore, performed current-clamp recordings in the same motoneurons as above. Starting from its natural resting membrane potential, we injected a triangular ramp of current to elicit the repetitive firing of the motoneuron (Figure 2A-B), and measured the voltage threshold for spiking on the first spike triggered by the ramp. In concordance with the hyperpolarization of the PICs, spiking threshold was also hyperpolarized in young animals (WT: −45.35±5.48 mV, N=19 vs. mSOD1: −54.86±5.15 mV, N=14; *g*=−1.74 95%CI[−2.53-−0.81]; *t*(31)=5.10, *p*=1.9e-05), and increased over time (Figure 1G).

**Figure 2.**
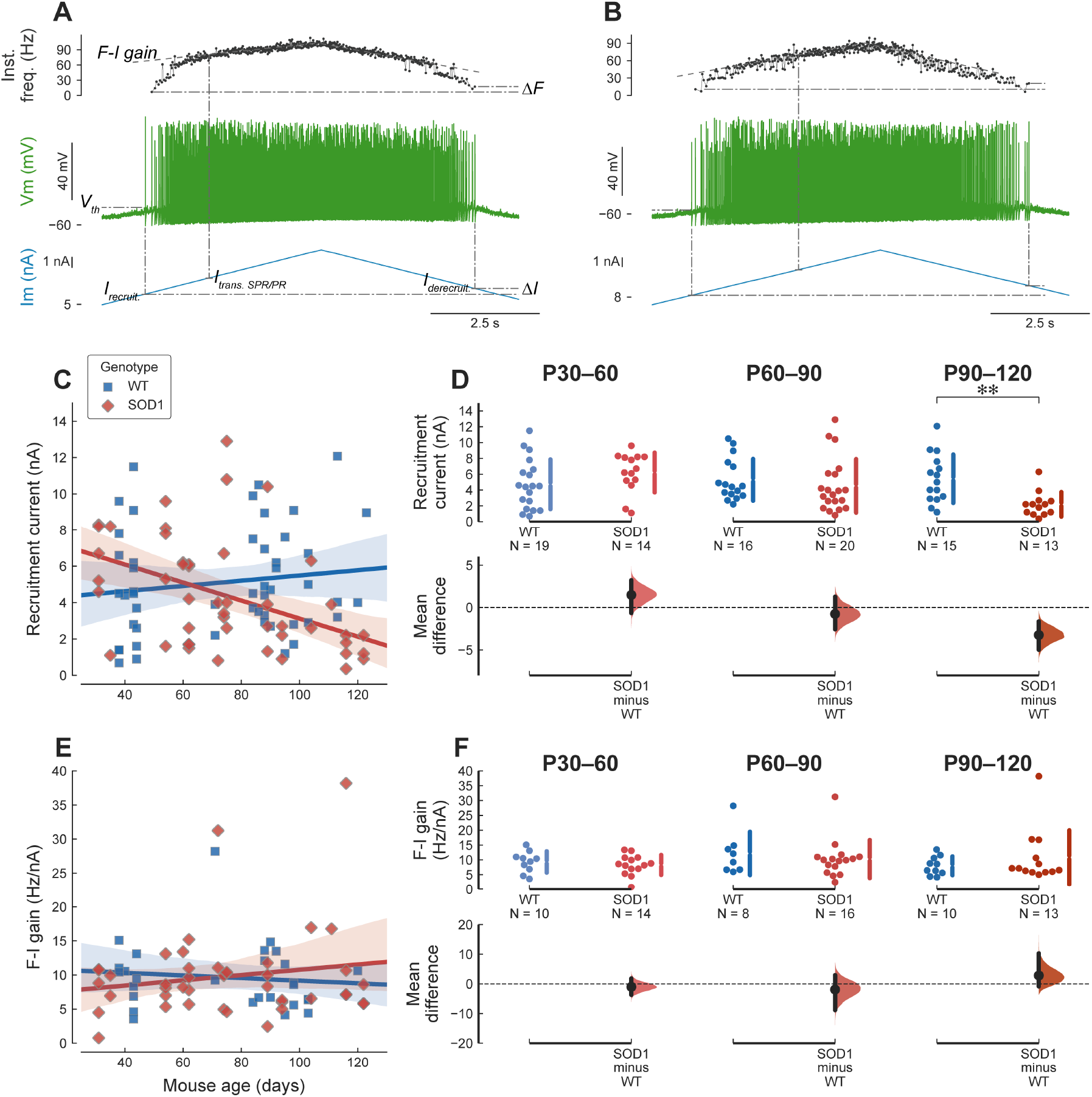
Mutant motoneurons are not hyperexcitable. **A**. Example of the response of a P43 WT mouse (same motoneuron as in Figure 1A) to a triangular ramp of current. From bottom to top, traces are: injected current, membrane potential, and instantaneous firing frequency. The dash-dotted lines show some of the measurements: *I_recruit_*.: current intensity at which the motoneuron starts to fire; *I_derecruit_*.: de-recruitment current; *ΔI*: difference between the de-recruitment and recruitment currents; *I_trans. SPR/PR_*: the current at the transition between sub-primary range and primary range; *F-I gain*: slope of the linear fit of the firing frequency in the primary range; *ΔF*: difference between the instantaneous firing frequency at de-recruitment and recruitment; *V_th_*: voltage threshold for spiking measured on the first spike of the ramp. **B**. Example of the response of a P35 mSOD1 mouse (same motoneuron as in Figure 1B). Same organization as in A. **C**. Plot of the current intensity required for eliciting the first spike on a triangular ramp of current in WT (blue squares) and mSOD1 motoneurons (red diamonds). WT: slope=0.1 nA/week 95%CI[−0.11-0.31], *r^2^*=0.019 (*p*=0.34). SOD1: slope=−0.35 nA/week 95%CI[−0.56–−0.14], *r^2^=0.2* (*p*=0.0019). **D**. Breakdown of the difference in recruitment current between WT and mSOD1 motoneurons in each of the age groups. In young adult and presymptomatic mice, mutant motoneurons require the same amount of current than WT motoneuron to fire, P30-60 WT: 4.73±3.14 nA, N=19 vs. mSOD1: 6.21±2.51 nA, N=14; *g*=0.50 95%CI[−0.24-1.24]; *t*(31)=−1.51, *p*=0.14. P60-90 WT: 5.28±2.64 nA, N=16 vs. mSOD1: 4.53±3.38 nA, N=20;, *g*=−0.24 95%CI[−0.93-0.41]; *t*(34)=0.75, *p*=0.46. At the symptomatic stages (P90-120), mutant motoneurons exhibit a lower current threshold for firing (WT: 5.43±3.07 nA, N=15 vs. mSOD1: 2.19±1.56 nA, N=13; *g*=−1.26 95%CI[−1.95-−0.47]; *t*(26)=3.59, *p*=0.0017), compatible with the loss of the least excitable cells. **E**. The slope of the F-I relationship, measured over the primary range, is not affected by the mutation, regardless of the age of the animals. WT: slope=−0.13 Hz/nA/week 95%CI[−0.65-0.38], *r^2^*=0.011 (*p*=0.6). SOD1: slope=0.27 Hz/nA/week 95%CI[−0.23-0.77], *r^2^*=0.028 (*p*=0.28). **F**. Breakdown of the difference between WT and mSOD1 motoneurons by age group: P30-60 WT: 9.3±3.6 Hz/nA, N=10 vs. mSOD1: 8.3±3.4 Hz/nA, N=14; *g*=−0.28 95%CI[−1.10-0.64]; *t*(22)=0.69, *p*=0.5. P60-90 WT: 12.2±7.3 Hz/nA, N=8 vs. mSOD1: 10.3±6.4 Hz/nA, N=16; *g*=−0.27 95%CI[−1.22-0.54]; *t*(22)=0.62, *p*=0.54. P90-120 WT: 8.0±3.2 Hz/nA, N=10 vs. mSOD1: 10.8±9.1 Hz/nA, N=13; *g*=0.38 95%CI[−0.43-0.93]; *t*(21)=−1.04, *p*=0.31.

### Young adult motoneurons are nonetheless not hyperexcitable

Despite the large alterations in PIC amplitude and activation voltage and the change in the voltage threshold for spiking, the excitability of the motoneurons was remarkably unaffected by the SOD1 mutation, regardless of age. Excitability was quantified using the intensity of the current required to elicit the first spike on the ascending ramp (“recruitment current”). Although the recruitment current decreased over time in mSOD1 animals (Figure 2C), this effect was mostly driven by older animals (Figure 2D). Young adult (P30-60) mutant motoneurons, whose neuromuscular junctions are just starting to be denervated, and pre-symptomatic motoneurons (P60-90), which experience substantial denervation, did not require, on average, less current to reach firing threshold than WT controls (Figure 2D). At symptomatic stages (P90-120), the SOD1 mutation does lead to a decrease in the recruitment current compared to WT animals (Figure 2D), which is probably caused by the degeneration of the high threshold motoneurons at this stage.

Neuronal excitability depends not only on how much current is needed to start firing, but also at what frequency the neuron is firing once it is recruited. We quantified the firing frequency of the motoneurons by measuring the slope of the frequency-current relationship. As shown previously (Manuel et al., 2009), most F-I curves, regardless of age and genotype, showed a distinct sub-primary range (SPR) with a steep slope and high variability followed by a linear phase called primary range (PR). We used the “gain” of the motoneuron, i.e. the slope of the F-I curve in the PR, as another measure of excitability of motoneurons. Despite the alterations in PICs in mutant motoneurons, the gain of the motoneuron F-I curves was unaffected by both age and mutation (Figure 2E-F).

### Compensatory changes responsible to maintain excitability

The fact that young adult motoneurons are not hyperexcitable despite substantial alterations in the PICs suggests that other compensatory mechanisms are acting to preserve the functional output of the cells. Motoneuron input conductance is an important factor controlling neuron excitability. We estimated the input conductance of mSOD1 and WT motoneurons from the slopes of their I-V relationships around the level of the resting membrane potential (see Figure 1 and Methods). Input conductances of mSOD1 and WT motoneurons are plotted against the age of the animal in Figure 3A. Consistent with the continuous increase in the size of the animals over the age span studied, WT motoneurons exhibit a moderate increase in input conductance in WT motoneurons over time (Figure 3A). On the other hand, mSOD1 motoneurons showed the opposite trend (Figure 3A). Young adult mutant motoneurons had an input conductance almost twice as high as WT controls at P30-P60 (Figure 3B). This increase is relatively greater (~1.7 fold) than that observed in neonatal motoneurons (~1.25 fold) (Quinlan et al., 2011), suggesting that the trend for increased conductance has become stronger as the animal matures into the young adult state. However, by the late pre-symptomatic stage (P60-90), the mutation had no longer any effect on input conductance. Finally, symptomatic (P90-120) mSOD1 motoneurons had a smaller input conductance than WT controls (Figure 3B), but this difference could be due to the degeneration of the largest, high-threshold units at this stage.

**Figure 3:**
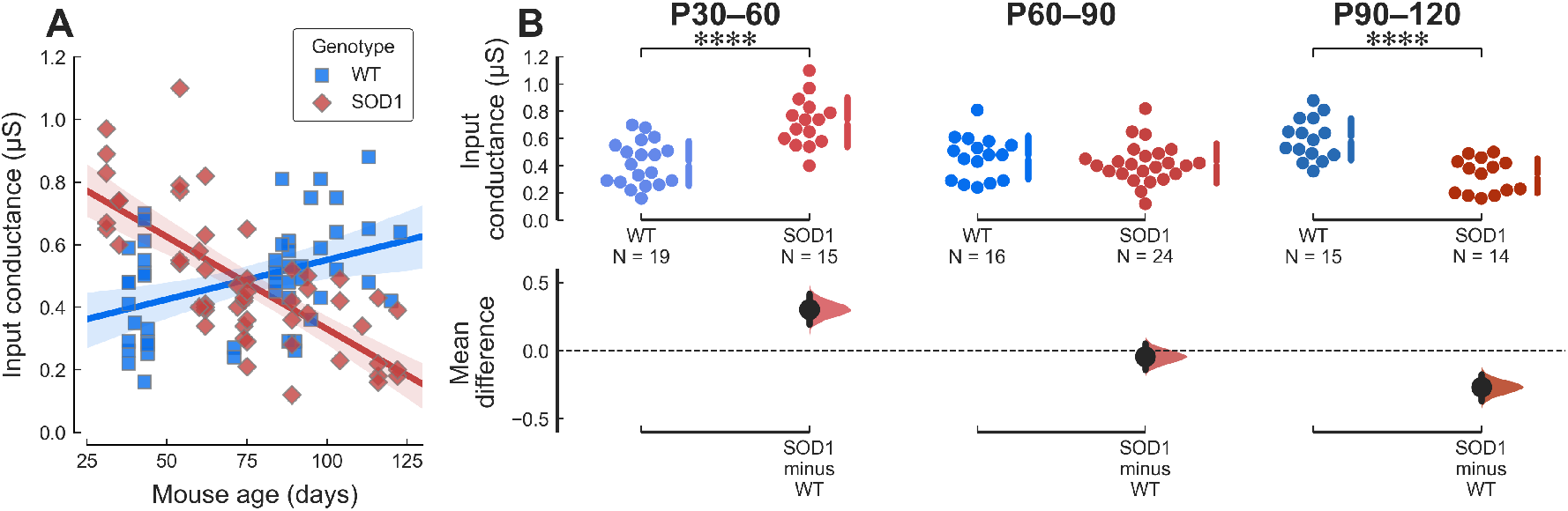
Young mutant motoneurons have an aberrantly large input conductance. **A**. Plot of the motoneuron input conductance vs. age in WT (blue squares) and mSOD1 (red diamonds) animals. The solid lines correspond to the linear regression lines with 95% confidence intervals (shaded areas). WT slope=0.018 μS/week 95%CI[0.0061-0.029], *r^2^*=0.16 (*p*=0.0035); SOD1 slope=−0.041 μS/week 95%CI[−0.052–−0.03], *r^2^*=0.53 (*p*=6.1e-10).) **B**. Breakdown of the difference in input conductance between WT and mSOD1 animals by age groups. P30-60 WT: 0.42±0.16 μS, N=19 vs. mSOD1: 0.72±0.18 μS, N=15; *g*=1.73 95%CI[0.92-2.41]; *t*(32)=−5.06, *p*=2.2e-05. P60-90 WT: 0.46±0.16 μS, N=16 vs. mSOD1: 0.42±0.15 μS, N=24; *g*=−0.28 95%CI[−0.96-0.37]; *t*(38)=0.88, *p*=0.39. P90-120 WT: 0.60±0.15 μS, N=15 vs. mSOD1: 0.33±0.13 μS, N=14; *g*=−1.85 95%CI[−2.57-−1.16]; *t*(27)=5.16, *p*=2.1e-05.

The increase in input conductance is not the only compensatory change happening in young adult mutant motoneurons. In particular, one would have expected the leftward shift in the activation voltage of PICs seen in these motoneurons to have a profound impact on their firing behavior. This discrepancy can be explained by a parallel hyperpolarization (by almost 10 mV) of the resting membrane potential in young adult mSOD1 motoneurons (Figure 4, P30-60). This effect is still observable at pre-symptomatic stages, but to a lesser degree (Figure 4, P60-90). At symptomatic ages, however, mutant motoneurons had similar resting membrane potentials as WT controls (Figure 4, P90-120).

**Figure 4.**
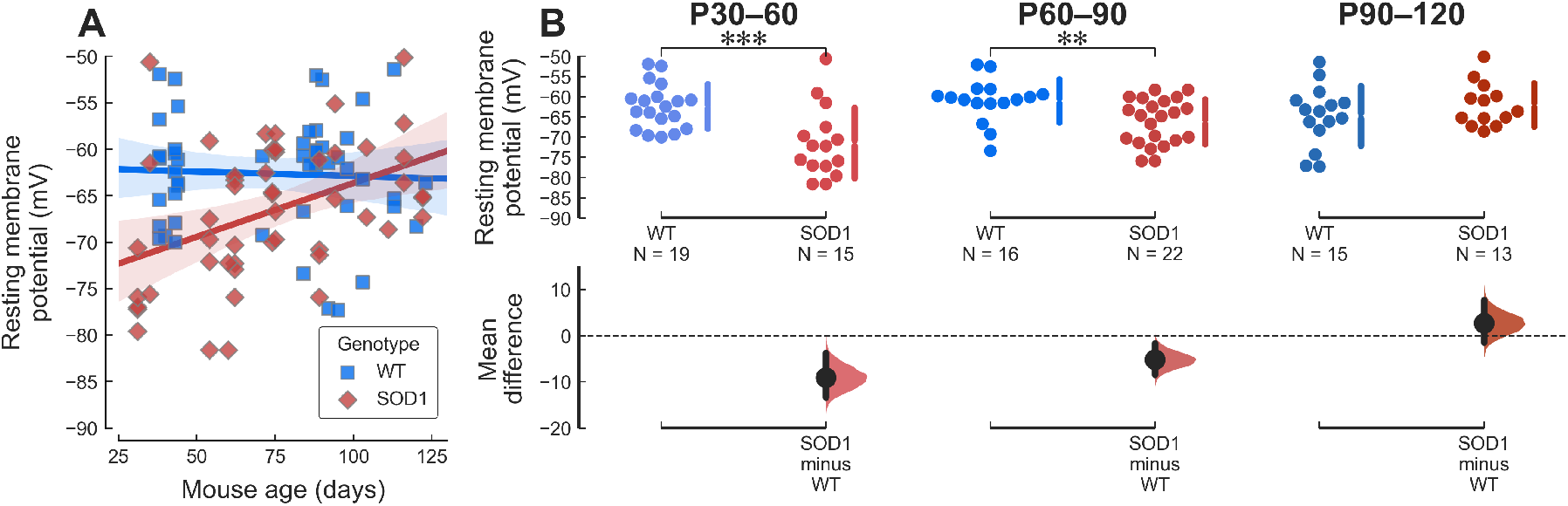
The resting membrane potential of young adult mutant mice is hyperpolarized. **A**. Plot of the motoneuron resting membrane potential vs. age in WT (blue squares) and mSOD1 (red diamonds) animals. The solid lines correspond to the linear regression lines with 95% confidence intervals (shaded areas). WT: slope=−0.068 mV/week 95%CI[−0.52-0.38], *r^2^*=0.0019 (*p*=0.76). SOD1: slope=0.81 mV/week 95%CI[0.31-1.3], *r^2^*=0.18 (*p*=0.0022). **B**. Breakdown of the difference in resting membrane potential between WT and mSOD1 animals by age groups. P30-60 WT: −62.38±5.52 mV, N=19 vs. mSOD1: −71.45±8.78 mV, N=15; *g*=−1.24 95%CI[−2.13-−0.23]; *t*(32)=3.49, *p*=0.002. P60-90 WT: −60.98±5.35 mV, N=16 vs. mSOD1: −66.18±5.63 mV, N=22; *g*=−0.92 95%CI[−1.54-−0.18]; *t*(36)=2.89, *p*=0.0067. P90-120 WT: −64.71±7.42 mV, N=15 vs. mSOD1: −61.99±5.43 mV, N=13; *g*=0.40 95%CI[−0.35-1.10]; *t*(26)=−1.12, *p*=0.27.

When taking into account this hyperpolarization of the resting membrane potential, the relative values of the voltage threshold for spiking (ΔV_th_, difference between the voltage threshold and the resting membrane potential), as well as the relative activation voltage of the PICs (ΔV_PIC_) and the relative voltage at the peak of the PIC (ΔV_peak_) were all similar between WT and mSOD1 motoneuron regardless of age (Table 1), which suggest that these are the quantities that are homeostatically regulated.

### Arms race between PICs and input conductance

Our results show that, during the disease progression, motoneurons are actively engaged in a homeostatic process to maintain their firing output. The ratio of PIC to conductance is a major determinant of net excitability. If these two parameters grow in proportion, then net excitability is likely to stay about the same (Huh et al., 2017). We thus analyzed the relationship between PIC amplitude and input conductance (Figure 5). In the young adult animals (Figure 5A), the majority of the mutant motoneurons are clustered in the upper right-hand corner, with values of both conductances and PIC amplitudes that are outside of the range of WT controls. Yet, the slopes were similar for both groups (ANCOVA, no significant interaction between input conductance and genotype on PIC amplitude, *t*(30)=−0.590, *p*=0.560), and the mutation had only a negligible effect on the relationship between PIC amplitude and conductance (ANCOVA, effect of mutation −0.40 95%CI[−6.73-5.92], *t*(30)=−0.130, *p*=0.898). Therefore the relationship between PIC amplitude and input conductance was the same in WT and mSOD1 mice. This implies that, in these animals, the cells that are abnormally large (large input conductance) also have PICs that have increased in proportion. It is plausible that this process would, by itself, cause undue stress to the cell (Attwell and Laughlin, 2001; Herculano-Houzel, 2011; Howarth et al., 2012), due to the high metabolic demand imposed by their large size, as well as the potential massive influx of calcium caused by their abnormally large PICs. At the pre-symptomatic stages, the WT and mSOD1 populations roughly overlap (Figure 5B). At the symptomatic ages, the situation is reversed, with the appearance of a cluster (N=6) of very small cells with very small PICs (Figure 5C). These results show that the overall initial increase in PIC and conductance amplitudes were driven by the SOD1 mutation generates a sub-population of motoneurons with conductance and PIC amplitudes that are well above the normal, WT range (P30-60). Then, as the disease progresses, the steady reduction in these amplitudes is so strong that by P90-120, some of the surviving cells fall well below the normal range.

**Figure 5.**
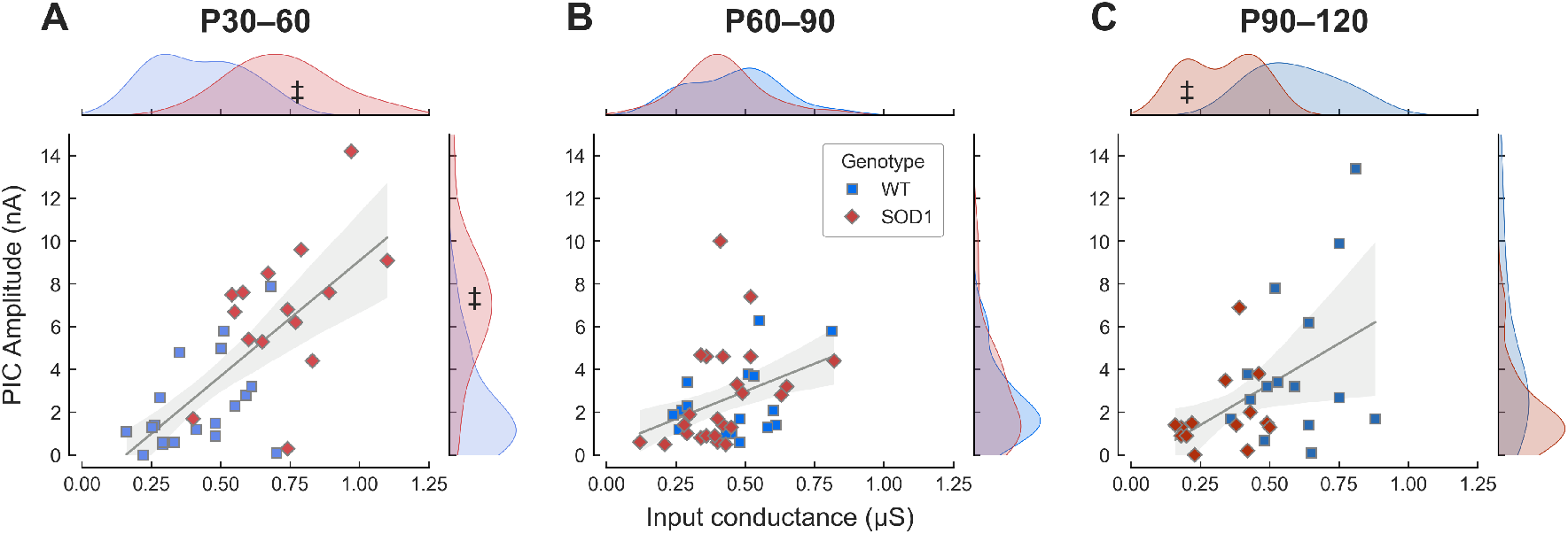
Some motoneurons exhibit properties outside of the normal range. **A**. Plot of the PIC amplitude vs. the input conductance of young adult motoneurons (P30-60) in WT (blue squares) and mSOD1 (red diamonds) animals. The grey line is the best linear fit ± 95% confidence interval (shaded area) for both samples. Slope=7.7 mV 95%CI[2.6-12.7], *r^2^*=0.55 (*p*=0.004). The marginal plots indicate the kernel density estimation of the distributions of the values in the two populations. The ‡ symbol points to the fraction of the mSOD1 population that is outside the range of the WT population. **B**. Same as A for the presymptomatic age range P60-90. Slope=5.4 mV 95%CI[1.1-9.8], *r^2^*=0.15 (*p*=0.016). **C**. Same as A for the symptomatic age range P90-120. Slope=7.0 mV 95%CI[−0.8-14.9], *r^2^*=0.23 (*p*=0.078).

### Some cells are hypoexcitable and cannot fire repetitively

Despite the remarkable ability of spinal motoneurons to maintain their excitability and firing output demonstrated above, some mSOD1 motoneuron tended to lose their ability to fire repetitively as the disease progressed (Delestrée et al., 2014; Martínez-Silva et al., 2018). Out of the 53 mutant motoneurons recorded, 6 motoneurons could not fire repetitively to current ramps despite still being able to fire a single or a few action potentials to current steps. Figure 6 shows an example of such a motoneuron. Although those cells were not, on average, larger than those that fired repetitively (Figure 6D), non-firing cells were characterized by a very small PIC amplitude (Figure 6E, see also (Huh et al., 2017)). Finally, although non-firing motoneurons could be recorded in WT mice (7 out of 50 motoneurons), non-firing motoneurons appeared, on average, 25 days earlier in mSOD1 mice (Figure 6F).

**Figure 6:**
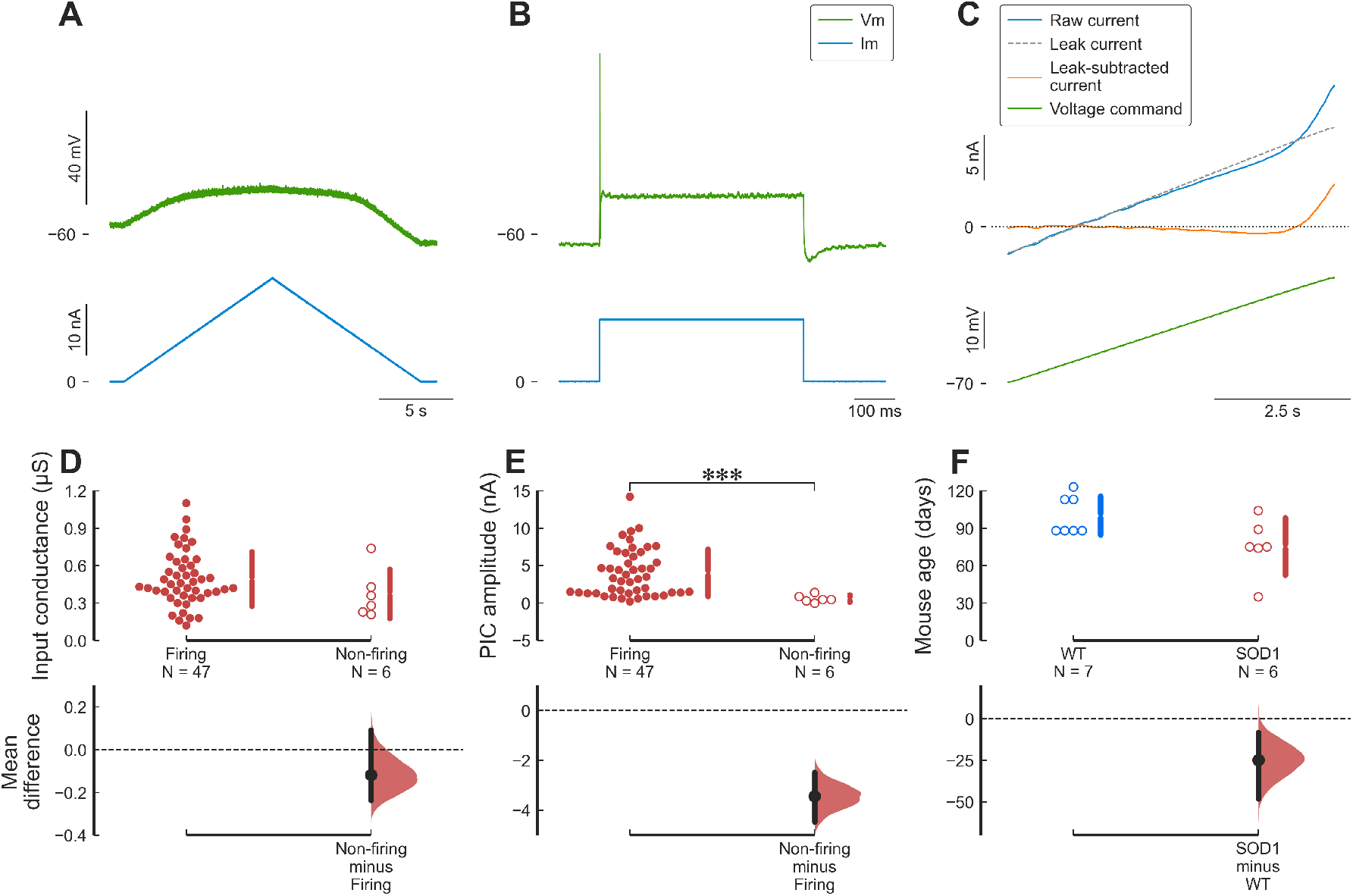
Some cells are hypoexcitable and cannot fire repetitively. **A**. Example of an mSOD1 motoneuron (from a P88 mouse) that is unable to fire repetitively in response to a triangular ramp of current. Top trace: membrane potential, bottom trace: injected current. **B**. This same motoneuron was nevertheless able to generate a single full-height action potential in response to a square pulse of current. Same organization as in A. **C**. Voltage-clamp measurement of the PICs in this same motoneuron. Traces are (from top to bottom), leak current (dashed line), raw current (blue), leak-subtracted current (red) and voltage command (green). **D**. Non-firing motoneurons had a similar input conductance compared to motoneurons capable of firing repetitively. Firing: 0.49±0.22 μS, N=47 vs. non-firing: 0.38±0.20 μS, N=6; *g*=−0.54 95%CI[−1.11-0.50]; *t*(51)=1.36, *p*=0.22 **E**. Non-firing motoneurons had much smaller PICs than motoneurons able to fire repetitively. Firing: Firing: 4.04±3.18 nA, N=47 vs. non-firing: 0.60±0.49 nA, N=6; *g*=−1.12 95%CI[−1.42–−0.86]; *t*(51)=6.81, *p*=1.3e-08 **F**. Non-firing motoneurons appear earlier in mSOD1 animals compared to WT animals. WT: 100±16 days old, N=7 vs. SOD1: 75±23 days old, N=6; *g*=−1.20 95%CI[−2.06-0.01]; *t*(11)=2.24, *p*=0.053.

### Other motoneuron properties

Given the large number of electrophysiological parameters measured in each cell (21 per cell), we used Principal Component Analysis (PCA) to analyze the overall behavior of the cells across time and genotype. The first three principal components (PCs) accounted for 67% of the variance in the data (PC1: 33.8%, PC2: 21.4%, PC3: 11.9%). Figure 7A_1-3_ shows how the first three principal components varied with the age of the animal in mSOD1 and WT mice. For WT mice, all principal components were constant over time (slopes: PC1 0.102 AU/week 95%CI[−0.178-0.381], *r^2^*=0.024, *p*=0.459; PC2 −0.112 AU/week 95%CI[−0.368-0.144], *r^2^*=0.034, *p*=0.376; PC3 0.043 AU/week 95%CI[−0.182-0.269], *r^2^*=0.007, *p*=0.694). On the other hand, in mSOD1 mice, the first two principal components evolved over time. PC1 started at a higher value than WT mice, decreased over time (slope −0.460 AU/week 95%CI[−0.625–−0.295], *r^2^*=0.450, *p*=1.6e-06), and became smaller than in WT mice at endstage. PC2 followed the opposite trend. In young mSOD1 animals, PC2 was lower than in WT animals, and then it increased over time (albeit by a very small amount; 0.151 AU/week 95%CI[0.006-0.296], *r^2^*=0.102, *p*=0.042). Finally, PC3 stayed constant (−0.089 AU/week 95%CI[−0.192-0.013], *r^2^*=0.074, *p*=0.086) and indistinguishable from WT. Higher principal components did not show any dependency on age or genotype (not shown).

**Figure 7.**
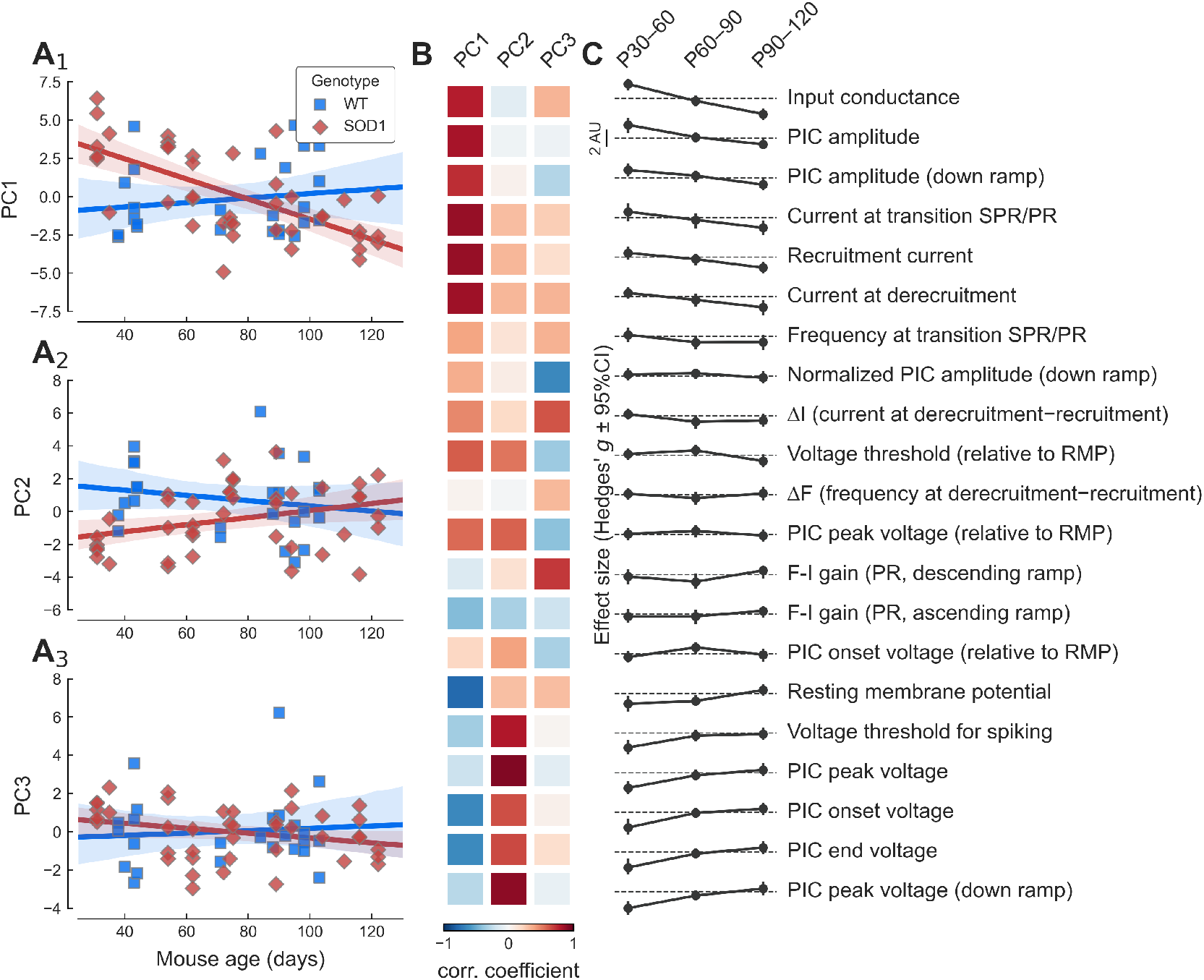
Overview of the change in principal components over time. **A_1-3_**. Plot of the three first principal components vs. age in WT (blue squares) and mSOD1 motoneurons (red diamonds). The solid lines correspond to the linear regression lines with 95% confidence intervals (shaded areas). **B**. Heatmap showing the correlation coefficient between each principal component (columns) and the features (rows) shown on the right. Correlation coefficients are color-coded from dark blue (*r*=−1) to dark red (*r*=+1). **C**. Summary of the evolution of the difference between WT and mSOD1 motoneurons (quantified by the effect size Hedges’*g*) for each of the features and each of the time points considered. The features are ordered by the size of the effect of the mutation in the P30-60 age group. On each row, the dashed line represents an effect size of zero. Scale bar: 2 units.

The opposite behavior of PC1 and PC2 in mSOD1 mice indicates that several features are anti-correlated. PC1 was strongly positively correlated to input conductance, as well as most current measurements (PIC amplitude, recruitment current, etc.) (Figure 7B), which were also the features that were increased the most in young adult mSOD1 vs. WT animals (Figure 7C, top rows). On the other hand, PC2 was more strongly correlated to voltage measurements (voltage threshold, PIC peak voltage, PIC onset voltage, resting membrane potential; Figure 7B), which were features that were strongly decreased in young adult mSOD1 vs. WT (Figure 7C, bottom rows). Overall, PCA clearly highlights two sets of features that evolve in opposite direction over time, presumably to compensate for one another in order to maintain neuronal excitability as close to normal as possible.

## Discussion

In this paper, we studied how motoneuron electrical properties evolve over the time course of ALS, focusing particularly on the persistent inward currents (PICs). We show that, in young adult mutant mice, before and up to the time when motor unit denervation is just starting, PICs are abnormally large compared to controls. This increase in PIC amplitude is accompanied by a parallel increase in motoneuron input conductance with the net effect that the excitability of the cells remains normal. Later, while the animals remain pre-symptomatic but denervation has begun, motoneuron properties return to normal levels. Finally, in symptomatic animals, mutant motoneurons tend to have smaller input conductance and smaller recruitment current, which is most likely due to the death of the largest, low threshold cells at this stage.

### Homeostatic regulation of motoneuron output

Although ALS is classically considered an adult-onset disease, we know from earlier studies of motoneurons in animal models of ALS that the change in motoneurons’ intrinsic excitability is the first sign to be seen in the pathogenesis, long before any overt motor deficits manifest. Hyperexcitability is observed in cultured E13 spinal motoneurons (Kuo et al., 2004), E15 cortical motoneurons (Pieri et al., 2009), E17.5 spinal motoneurons (Martin et al., 2013 p.20), as well as early postnatal hypoglossal motoneurons (van Zundert et al., 2008). Consequently, the motor system in general, and motoneurons in particular, must exhibit remarkable capability for homeostatic regulation to maintain a quasi-normal motor output until overt symptoms appear, but there are many combinations of intrinsic properties that can produce the same firing pattern (Marder and Goaillard, 2006). In neonatal mSOD1 mice, the majority of motoneurons exhibited normal excitability (based on recruitment current and F-I gain), although the most resistant population of motoneuron seem to retain some hyperexcitability (Quinlan et al., 2011; Leroy et al., 2014). Yet, PICs were almost twice as large in mutant compared to WT neonatal motoneurons (Quinlan et al., 2011). This increased PIC amplitude was seemingly compensated by a parallel increase in input conductance (Quinlan et al., 2011). In adults, this trend seems to continue, particularly in the young adult age group. Global analysis of our dataset using PCA shows that, generally speaking, currents tend to be much bigger in young adult mSOD1 mice compared to WT (we observe an almost 3× increase in PIC amplitude, accompanied by an almost 2× increase in input conductance), which then tend to decrease over time. This increase is accompanied by a hyperpolarization of the PIC onset and peak voltages, but which are compensated by a commensurate hyperpolarization of the resting membrane potential. These trends with disease progression from the embryonic state through the development of severe symptoms are summarized in Figure 8. An overall pattern of oscillations is evident, with the PIC oscillations tending to increase excitability but the conductance and rest potential oscillations tending to reduce it.

**Figure 8:**
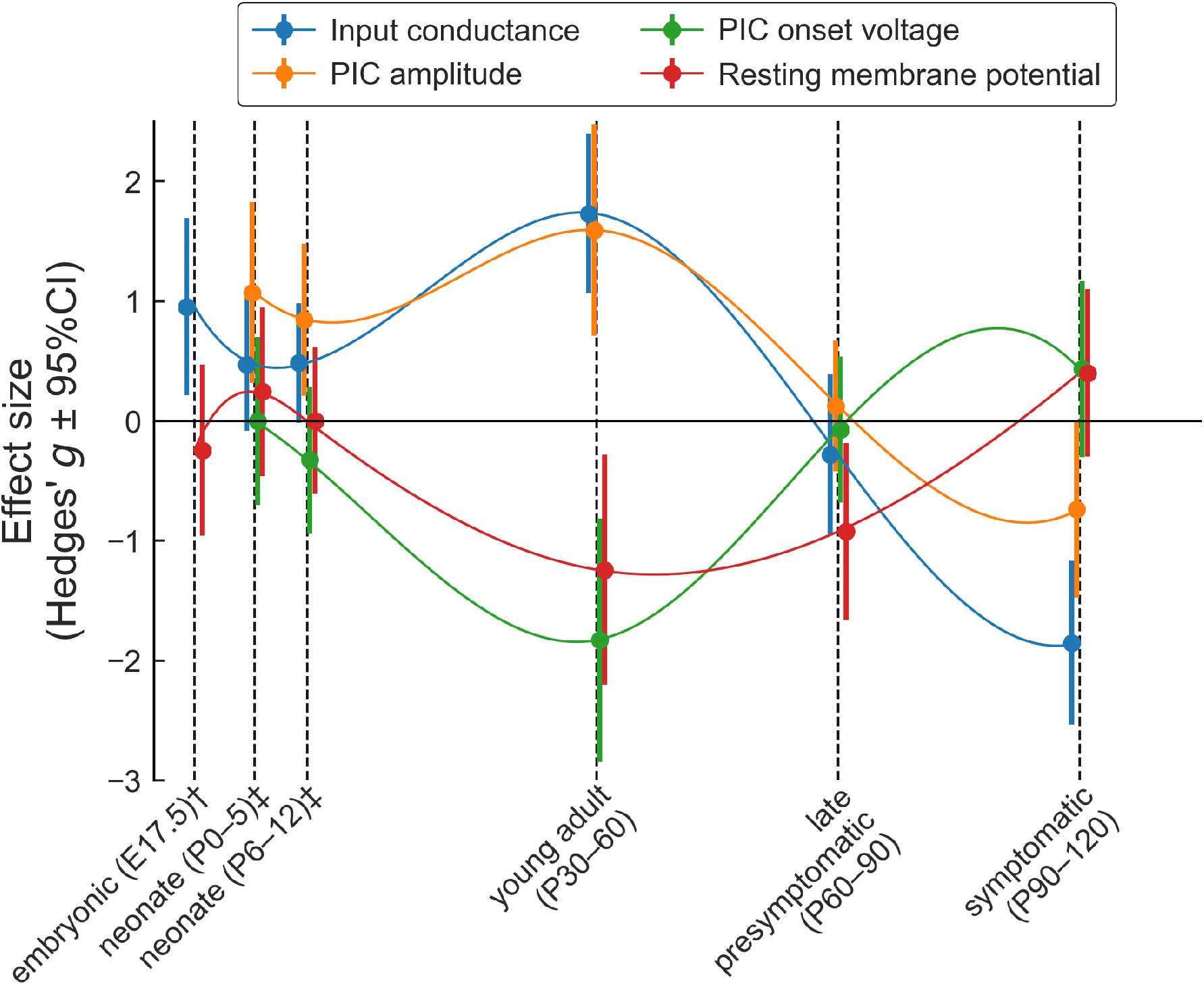
summary of the changes in motoneuron properties over time. Schematic representation of the changes in four key electrophysiological properties over time. The dots represent the effect size (Hedges’ *g*) and the vertical bars show the 95%CI around *g*. The thin lines are cubic splines interpolation of the data over time. The points have been slightly staggered so that the vertical bars do not occlude each other. †Data from embryonic motoneurons are from Martin et al. (2013). These authors did not measure PICs in embryonic motoneurons. Kuo et al. (2004) did measure PICs, but their embryonic motoneurons were cultured for 10-30 days in vitro, and their development stage is therefore uncertain. ‡Data from neonates (P0-5 and P6-12) are from Quinlan et al. (2011).

The properties of the motoneurons seem to normalize at the late pre-symptomatic stage (P60-90), but it is unclear whether this is due to changes in the homeostatic pathways involved, or whether it is caused by the start of the degeneration process in the most vulnerable motoneurons. At symptomatic stages, motoneurons appear to be hyper-excitable. They have, on average, a smaller input conductance (Figure 3), and smaller recruitment current (Figure 2), but that most likely reflect the fact that the largest cells have degenerated by this age, and that the remaining cells may have shrunk (see below, and Dukkipati et al., 2018). In addition, it should be noted that a small but growing number of cells become incapable of firing repetitively at this stage (Delestrée et al., 2014; Martínez-Silva et al., 2018).

### The hidden cost of excitability homeostasis

Although the degeneration of motoneurons in ALS constitutes a failure in cell homeostasis, our results show that, throughout the “silent”, presymptomatic phase of ALS, motoneurons are actively engaged in a highly successful homeostatic process to maintain their firing output, and thereby producing a normal motor behavior.

The driving force behind this homeostatic process remains mysterious. Although it is commonly agreed that intracellular calcium plays a major role in this process (Turrigiano et al., 1994; O’Leary et al., 2010; O’Leary and Wyllie, 2011), this calcium can enter the cell through various channels (e.g. synaptic receptors, voltage-dependent calcium channels) and the location and pattern of that calcium signal could potentially have distinct downstream targets. We demonstrated here that motoneurons can maintain a normal firing output through most of the pre-symptomatic phase of the disease in response to current injected through the microelectrode. Yet, during behavior, motoneurons are activated through synaptic inputs, most of which are impinging on the dendritic tree. Bączyk, Ouali Alami et al. (2020) have recently demonstrated that excitatory synaptic inputs are depressed in pre-symptomatic mSOD1 mice. Excitability homeostasis, therefore, cannot be restricted to the intrinsic properties of motoneurons, but potentially involve the whole sensorimotor network. Indeed, in the motor cortex, although several cell types (including corticospinal pyramidal cells) were found to be intrinsically hyperexcitable in symptomatic mSOD1 mice, the activity of corticospinal pyramidal neurons during behavior (measured by two-photon calcium imaging during head-fixed locomotion) was indistinguishable from controls (Kim et al., 2017). Whether network-level homeostasis and intrinsic excitability homeostasis are parallel, independent, processes, or whether one precedes and drives the other remains to be investigated.

Whatever the case may be, the process by which the cells maintain the firing output could, by itself, be a major source of stress for the cells. We do not know whether the cells increase their conductance to compensate for the fast increase in PICs or *vice versa*, but the result is the appearance of cells with both input conductance and PIC amplitudes outside of the normal range (Figure 5A). The increase in input conductance could be due to the insertion of more channels in the membrane. However, direct measurements of soma sizes have revealed that pre-symptomatic mutant cells are physically larger than controls (Shoenfeld et al., 2014; Dukkipati et al., 2018). This increase in size, coupled with the larger calcium entry in the cells due to the larger PICs (which are in part calcium-medicated (Li and Bennett, 2003)), is bound to cause undue metabolic stress on those cells that are outside of the normal range (Attwell and Laughlin, 2001), even though cells manage to maintain a normal output. If we assume that all motoneurons experience the same shift in their properties, then Fast Fatigable (FF) motoneurons, which are the cells with the largest input conductance, and the largest PICs in normal conditions (Lee and Heckman, 1998; Huh et al., 2017), are likely to be the cells that are the further out of the normal range. This might explain why they are the most vulnerable to ALS (Pun et al., 2006; Hegedus et al., 2007, 2008).

As the disease progresses, those large, vulnerable cells become the first motoneurons that will embark on a degeneration pathway (Saxena et al., 2009, 2013; Martínez-Silva et al., 2018), and will lose their ability to fire repetitively (Martínez-Silva et al., 2018). This early loss of the large cells is consistent with stress induced by aberrantly large conductances and PICs. This loss is also likely the primary reason why motoneurons appear to recover normal properties at late pre-symptomatic stages. Nevertheless, it is probable that all motoneurons, regardless of type, experience this form of stress. Interestingly, we observed that some of the smallest motoneurons, which are the most resistant to the disease, appear to shrink in the oldest animals (Figure 5C), consistent with anatomical observations (Kiernan and Hudson, 1991; Dukkipati et al., 2018). At this point, it is unclear whether this shrinkage is a pathological feature or a strong homeostatic effort by the surviving motoneurons to counteract the stress generated by their initial hypertrophy.

### Hyper-vigilant homeostasis

Mitchell and colleagues have recently performed meta-analyses of data obtained from many studies on multiple cellular properties, which revealed patterns of oscillations as the disease progressed, suggesting the existence of high feedback gains for homeostatic processes (Mitchell and Lee, 2012; Irvin et al., 2015). Several aspects of our results are consistent with this “hyper-vigilant” homeostasis hypothesis. Similar oscillatory patterns are present in our results, as illustrated in Figure 8. Although the cells are successful at maintaining their net excitability constant, this is achieved at the cost of excessively high amplitudes for conductances and PICs. These large amplitude changes lead to the appearance of groups of cells that are first well above the normal range of properties, followed by an over-reaction where some cells end up well below the normal range. This behavior fits well with the “hyper-vigilant” model (Irvin et al., 2015), but a direct test of the hyper-vigilant homeostasis hypothesis will require quantitative comparisons of the responses of ALS and WT motoneurons to controlled homeostatic challenges.

### Comparison with previous studies

Previous studies in adult SOD1 mice have suggested that spinal motoneurons became hyperexcitable (Meehan et al., 2010; Jensen et al., 2020). The most recent study uses the same SOD1(G93A) (albeit on a C57BL/6 genetic background) and same anesthetics as the present study, but has focused on two fairly advanced time points (~P75 and ~P115). They have observed an increase in input resistance (decrease in input conductance) between WT and mSOD1 mice at both P75 and P115. The increase at P115 is consistent with our results, and is probably due to the loss of the largest motoneurons at this stage. However, we did not observe a difference in resistance in our P60-90 group. The cause for this discrepancy remains unknown, but it could suggest an earlier onset of cell death in their colony. In their dataset, this decrease in conductance was associated with an apparent hyperexcitability: the recruitment current was lower and the F-I curve was steeper. However, their use of a low DCC switching rate (3 kHz, compared to 6-8 kHz used here) may have distorted the firing properties of their cells and led to an overestimation of their excitability (Manuel, 2020 in press).

We have previously studied the properties of motoneurons in the same SOD1(G93A) mouse model of ALS (Delestrée et al., 2014; Martínez-Silva et al., 2018) but have mainly focused on a short period just at the onset of denervations (P45-55). Although that time point partly overlaps with the present young adult stage, we had not detected differences in input conductances (Martínez-Silva et al., 2018). This might be because the difference in input conductance is largest at earlier time points (Figure 3). Indeed, the difference is gone at P60-90, and we did report an increase in input conductance in an earlier series of experiments, which was more pronounced in younger animals (Delestrée et al., 2014). Similarly, we did not detect a difference in resting membrane potential in the Martínez-Silva et al. (2018) study. Again, the discrepancy could be due to the difference in the distributions of the ages of the animals studied. The precise reason for this difference remains, however, unknown and warrants further study.

Since input conductance is dependent on motoneuron size (Burke, 1981; Heckman and Enoka, 2012), our present results are consistent with anatomical studies that have measured soma sizes in mSOD1 mice. In young adult animals, motoneurons are markedly larger in mSOD1 mice compared to WT animals (particularly in males)(Shoenfeld et al., 2014; Dukkipati et al., 2018). At symptomatic stages, however, the situation is reversed and the remaining motoneurons appear to have shrunk below the size of the smallest motoneurons in WT mice (Dukkipati et al., 2018).

In the present study, we recorded only 13 motoneurons out of 103 (7/50 WT and 6/53 mSOD1) that were unable to fire repetitively, contrary to our previous study where we showed that a large proportion of vulnerable motoneurons became hypoexcitable before the onset of denervation (P45-55, Martínez-Silva et al., 2018). This discrepancy could be explained by the experimental constraints of the present study. Indeed, obtaining stable voltage-clamp recordings mice in vivo is quite a challenge, requiring electrodes able to pass substantial amounts of current and motoneuron able to withstand the protocol. Since we have only included for analysis cells in which both the voltage-clamp and current-clamp protocols were completed successfully, we hypothesize that we may have biased our sample towards the healthiest cells that have yet to take the path of degeneration.

### Limitations

The results presented here have been collected on the SOD1(G93A) mouse model of ALS, a model that has attracted criticisms due to it being an overexpression model, and its failure to bring seemingly promising therapies to the clinic (Philips and Rothstein, 2015). Nevertheless, some parallel with other studies suggests that our results could be extended to other models. First, Meehan et al. (2010) have studied an unrelated mSOD1(G127X) model and have shown that although the net excitability of the motoneurons was not affected by the mutation, there were nonetheless signs of increased PICs in these motoneurons. Second, we observed that, in late symptomatic animals, some cells exhibited a very small input conductance, well below the normal range at this age, suggesting that these cells actually shrank. This observation matches morphological measurements in mSOD1 mice (Dukkipati et al., 2018), but also observations from sporadic human patients (Kiernan and Hudson, 1991).

We demonstrated a remarkably successful homeostatic control of motoneuron firing, that we can attribute to commensurate changes in PICs and input conductance. However, it is likely that other currents are also implicated in this process (Marder and Goaillard, 2006). Limitations inherent to in vivo electrophysiology prevent us from isolating many different currents and further in vitro investigations in adults (Jiang and Heckman, 2006; Mitra and Brownstone, 2011; Jiang et al., 2017; Bhumbra and Beato, 2018) are warranted to shed more light on the panoply of channels involved in this process.

### Conclusion

Overall, our results show homeostasis for net excitability in mutant SOD1 motoneurons is remarkably strong in the presymptomatic state. This success however comes at a cost of large compensatory changes in basic electrical properties. These results support the hypothesis that ALS is not due to a single specific root cause but instead caused by an inherent instability at the system-level (Mitchell and Lee, 2012), possibly caused by a “hyper-vigilant” homeostatic system in motoneurons. Initial perturbations in the electrical properties of motoneurons would be overcompensated for, leading to new sources of stress, which, in turn, would be overcorrected, and so forth until the metabolic burden becomes too high to sustain for the motoneuron. FF motoneurons have higher metabolic needs (Le Masson et al., 2014) and lower calcium-buffering capabilities (Grosskreutz et al., 2010) in the first place, which make them particularly vulnerable to this vicious cycle.

## Acknowledgments

This work was supported by NIH NINDS R01NS077863 and NINDS R01NS110953. S.H. was supported by a National Science Foundation Graduate Research Fellowship (NSF GRFP).

## Data availability

Data files and a computational notebook allowing reproducing the analysis and figures of this article are provided at the URL: https://doi.org/10.5281/zenodo.3831946

